# Inference of maternal allele inheritance via Bayesian hierarchical model in noninvasive prenatal diagnosis

**DOI:** 10.1101/051995

**Authors:** Chunxuan Shao

## Abstract

Noninvasive prenatal diagnosis (NIPD) poses a promising solution for detecting genetic alterations in fetus genome. However, the inference of the maternal allele inheritance in monogenic autosomal recessive disease is still challenging. Here the Bayesian hierarchical model is proposed to deduce the allele inheritance basing on haplotype frequency. The Bayesian approach, which does not depend on the knowledge of fetus DNA proportion in maternal plasma, provides accurate estimations on both real and simulated data; moreover, it is most robust than current methods in analyzing noisy or even erroneous data.

## Introduction

The discovery of free fetal DNA in maternal plasma starts a new era of non-invasive prenatal diagnosis (NIPD).^1^ The placenta origin cell-free DNA could make up to 20% of total cell-free DNA in plasma, providing necessary materials for enormous applications.^2^ For example, sex determination by RT-PCR could achieve near 100% sensitivity and specificity.^3,4^ More recently, with the advent of next generation sequencing, NIPD of aneuploidy and subchromosomal abnormalities are feasible or even recommended for high risk patients in the clinical practices. ^5–7^

Combining the causative mutations identified in monogenic autosomal recessive diseases and deep sequencing, NIPD is of great value in clinical diagnoses and treatments via determining the parental alleles inheritance in fetus genome. NIPD has correctly revealed the zygosity of HBB in β-thalassemia families, and been applied in a study of 14 congenital adrenal hyperplasia families^9–10^. Although inferring the paternal alleles inheritance is relatively straight forward, it is still challenging to infer the inheritance of heterogeneous maternal alleles in recessive disease. Relied on the imbalance of maternal haplotypes in maternal plasma DNA mixed with fetus DNA, relative haplotype dosage analysis (RHDO)^8–10^, haplotype counting analysis^11^ and hidden Markov model (HMM)^12^ have been employed to dedue the maternal alleles inference. However, the proportion of fetus DNA in total cell-free DNA in mother plasma, which is a key parameter in above methods, might not be accurately estimated.

The RHDO analysis is based on the observation that the proportion of heterozygous maternal SNPs detected from mother blood is not different from 0.5 if there is no fetus DNA existed, and inherited maternal alleles from fetus makes the proportion shift away. The sequential probability ratio test (SPRT)^8,9^ is employed to perform the hypothesis testing with the null hypothesis states that the proportion of heterozygous maternal SNPs is equal to 0.5, via cumulatively adding the reads of SNPs from high-throughput sequencing data on each haplotype and comparing the ratio to a pre-defined threshold. The ratio and counts data used by SPRT recall a classical Bayesian example of inferring the bias of a mint by flipping coins and counting number of heads, where a hierarchical model has been shown to be a powerful approach. We found that the Bayesian hierarchical model correctly deduced the alleles inheritance in real and simulated data and provided the results in intuitive and informative ways. More importantly, we showed that the Bayesian approach is more robust in the context of noisy or even erroneous datasets.

## Methods

### Relative haplotype dosage analysis

Relative haplotype dosage analysis (RHDO), which makes use of high-throughput generation sequencing data of heterozygous maternal SNPs and homozygous paternal SNPs flanking interested alleles, aims to identify the subtle haplotype imbalance caused by the mixture of fetus DNA and thereafter deduces maternal alleles inheritance in fetus genome^9^. Briefly, type α and type β SNPs are selected from maternal haplotypes (HapI and HapII) based on their identity to corresponding paternal alleles, i.e., maternal SNPs in HapI that are same to the paternal SNPs are identified as type α SNPs, and maternal SNPs in HapI that are different to the paternal SNPs are identified as type β SNPs. Then the sequential probability ratio test (SPRT),^8,9^ which tests the haplotype imbalance of type α or β against predefined threshold based on the fetus DNA proportion, is used to determine if the proportion of either haplotype is significantly different from 0.5. This method has been extended to the scenarios where both maternal and paternal SNPs are heterozygous^9^.

### Bayesian hierarchical model

The bias *θ*_*s*_ is defined for each heterozygous maternal SNP as the proportion of reads mapped to HapI, i.e., counts_(HapI)_ / (counts_(Hap1)_ + counts_(HapII)_). The HapBias, *ω*, is defined as the proportion of HapI in mather plasma DNA. The Bayesian hierarchical model is depicted as:

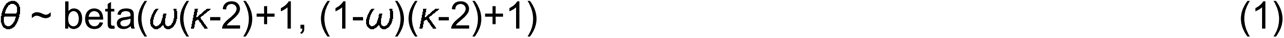

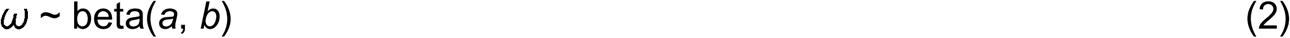

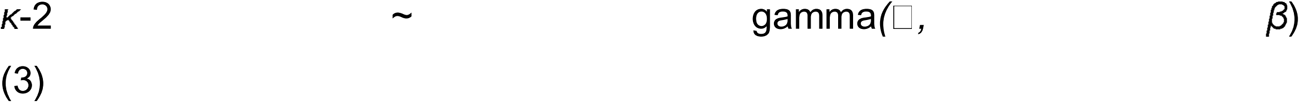

In formula (1), *θ*_*s*_ is modeled by a beta distribution using parameters of mode *ω* and concentration *κ*. The prior of *ω* is set to be a beta distribution using shape parameters *a* and *b*. *κ-2* is modeled by a gamma distribution using shape □ and rate *β*. The structure of Bayesian hierarchical model is summarized in Figure 1 in a bottom-up way. For each SNP, the read counts assigned to SNPs could be viewed as numbers of heads or tails generated in a series of Bernoulli process with a SNP specific bias *θ*_*s*_, which is modeled by a beta distribution with mode *ω* and concentration *κ*. The individual value of *θ*_*s*_ will be near to *ω*, and *κ* controls how close *θ*_*s*_ is to *ω*. We set the prior distribution of *ω* to be beta(2, 2) to reflect the fact that the HapBias is close to 0.5, and let the data play a major role in determining the posterior distribution. We set the prior gamma distribution with mean of 10 and standard deviation of 100. The Markov chain Monte Carlo simulation is used to approximate the posterior distribution of *θ*_*s*_, *K* and *ω*.

**Figure 1.**
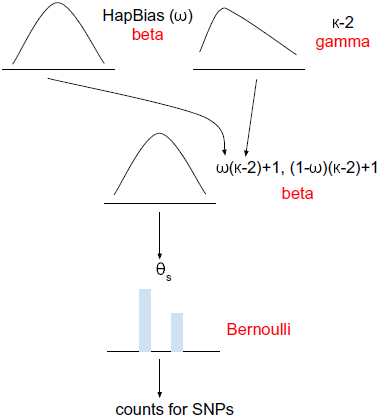
A scheme describes the Bayesian hierarchical model used to infer maternal alleles inheritance. The read counts are assumed to be generated in a Bernoulli process with a SNP specific bias *θ*_*s*_, which depends on a beta distribution with hyperparameter *ω* and *κ*. The parameter *ω,* which is referred as HapBias as well, has a prior beta distribution, whereas *κ* has prior gamma distribution. Names of probability distribution are highlighted with red.

Two outputs are possible for type α SNPs based analysis: HapBias is very close to 0.5 or HapBias is larger than 0.5, and the latter scenario indicates that HapI is overrepresented and hence inherited by the fetus. Similarly, two outputs are possible for type β based analysis: HapBias is very close to 0.5 or HapBias is smaller than 0.5, which indicates HapI is underrepresented and HapII is inherited by the fetus. The significance of over/under-represention of haplotype frequency is determined by the null hypothesis testing based on the relative location between high density interval (HDI) of the distribution of HapBias *ω* and region of practical equivalence (ROPE), which described in the next section.

### Null hypothesis testing in Bayesian analysis

In the procedure of the allele inheritance deducing, the null hypothesis is that the HapBias *ω* is equal to 0.5. In Bayesian analysis, a common approach to perform the null hypothesis testing is to examine the relative location between highest density interval of posterior distribution (HDI) and region of practical equivalence (ROPE).^13,14^ Parameter values inside HDI have higher probability density comparing with values outside this region, and sum to a defined probability (95% in our analysis).^13^ ROPE, or alternatively named indifference zone, is popularized by Freedman et al. as the counterpart to the single null value used in frequentist field.^15^ ROPE means the parameter values within this range are recognized to be practically equivalent for the null value. We denote (Δ_L_, Δ_H_) as the limits of HDI and (δ_L_, δ_H_) as the limits of ROPE. Six types of relation location between HDI and ROPE exist ((Fig. 2).^16^ We reject the null hypothesis that the HapBias is equal to 0.5 if there is no intersection between the HDI and the ROPE, and accept the null hypothesis if the ROPE completely contains the HDI.

**Figure 2.**
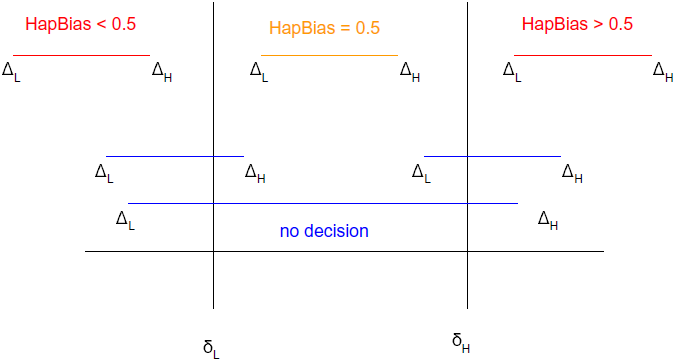
The Null Hypothesis testing based on ROPE and HDI. (ΔL, ΔH) and (δL, δH) are the limits of HDI and ROPE of 0.5, respectively. We reject the null hypothesis if there are no intersection between HDI and ROPE (highlighted in red). If ROPE contains HDI completely, we accept hypothesis that HapBias is equal to 0.5 (highlighted in orange). We make no decision for three other conditions (highlighted in blue).

### Limits of ROPE of the null hypothesis

The limits of ROPE are determined by practical purposes and overall considerations of experiments. In principle, wide ROPEs lead to the acceptance of null value, and vice versa for narrow ROPEs. We calculated the ROPE limits from the perspective of next-generation sequencing error. Assuming the error rate of sequencing platform is *e*, the total read counts for a single SNP on both HapI and HapII is *C*. In the null condition, the lower and upper limits of read counts of a SNP on HapI is *C/2 - C*e/2* and *C/2 + C*e/2,* respectively. Therefore, independent of the sequencing depth, the proportion of reads on HapI of a single SNP lies in the range [1/2 - e/2, 1/2 + e/2]. As the sample principle applies to all SNPs, the range is true for the haplotype as well. Thus, the range [1/2 - e/2, 1/2 + e/2] is used as the ROPE limits of HapBias of the null hypothesis.

### Simulation

We simulated the read counts data of haplotypes in the following steps: 1) we focused on the type α SNPs in the scenarios that the HapI was overrepresented; 2) the proportion of fetus DNA was set to be *p* (0.2%, 0.5%, 0.8%, 1%, 2%, 3%, 4%, 5%, 10%); 3) the sequencing depth C is set to be 200 for each SNP; 4) we used 500 SNPs and 1000 SNPs in the simulations, respectively; 5) the read counts for SNPs on HapI and HapII are (1+p)*C/2 and (1-p)*C/2, respectively; 6) we sampled data for each SNP with replacement to introduce the stochastic noise. We generated a dataset with p was set to be 0, indicating there is no fetus DNA in the maternal plasma.

## Results

### Application of the Bayesian hierarchical model on read data sets

We evaluated the Bayesian hierarchical model on high-throughout sequencing data to access its accuracy. The data were taken from a NIPD study with two β-thalassemia patient families where SNPs flanking interested genes were sequenced to generate read.^9^ To obtain the posterior distribution of HapBias, we performed 100,000 MCMC simulations with four chains (i.e., four individual simulation with difference initiation value). The ROPE of (0.49, 0.51) (the rationale will be discussed in a later section) and the 95% HDI of HapBias posterior distribution were used to perform null hypothesis testings. In the first family, the fetus inherited the HapII, and there are five type α and 46 type β SNPs available. The posterior distribution of HapBias using type β SNP demonstrated a 95% HDI region with limits of (0.435, 0.478), lied completely downstream of the ROPE, indicating HapI is underrepresented and HapII inherited in fetus, consistent with the publication results (Fig. 3A). On the other hand, the posterior distribution of HapBias based on type α SNPs showed that the ROPE completely fall within the HDI region, thus no conclusion could be drawn (Supplementary Fig. 1).

As Majority of analyzed SNPs in both parents in the second family are heterozygous, additional tagging SNPs were used to identify 24 type α SNPs.^9^ The Bayesian hierarchical model revealed that the ROPE lied completely upstream of 95% HDI, indicating the HapI was inherited in the fetus, consistent with known results (Fig. 3B) as well. Taken together, the Bayesian hierarchical model correctly deduced the maternal allele inheritance and represented the results in an intuitive way.

**Figure 3.**
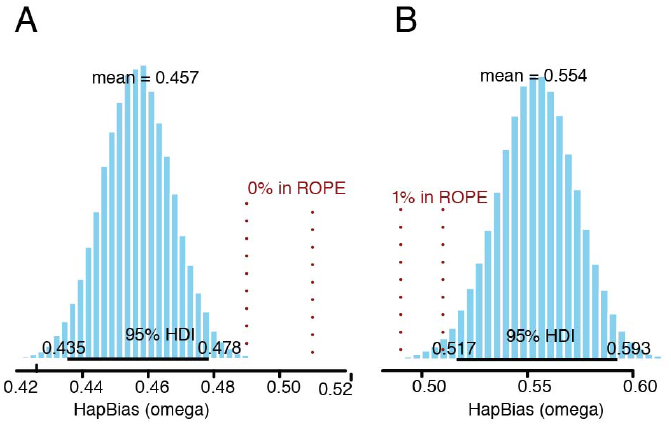
Posterior distribution of HapBias by the Bayesian hierarchical approach. (A) 95%HDI locates completely downstream of ROPE region for type β SNP in family one. (B) 95%HDI locates completely upstream of ROPE region for type a SNP in family two.

### Examination of Markov chain Monte Carlo simulation

It is important to check the quality of MCMC simulations used in the Bayesian model. We employed various visualization methods to examine two critical characters, convergence and accuracy. The trace plot, which shows the parameter in simulated chain steps, is useful to provide an overview of the convergence.^17,18^ The overlapping of trace plots suggested the MCMC simulations converged in both analyses (Fig. 4A, Supplementary Fig. 2A). Meanwhile, we summarized the parameter value sampled in individual chains as density plot. The overlapping of density plot suggests convergence of simulation as well (Fig. 4B, Supplementary Fig. 2B). The shrink factor, which reflects the ratio between between-chain variance and within-chain variance during simulation, is a generally used numerical measurement for convergence. The shrink factor is equal to one if all chains converged, and larger than one if orphan chains exist.^19^ Practically, the MCMC simulations with the shrink factor greater than 1.1 indicate lack of convergence.^13^ The mean and 97.5% quantile value of shrink factors gets close to 1 very quickly after the initial burn-in period in our data (Fig. 4C, Supplementary Fig. 2C), indicating the convergence is archived.

**Figure 4.**
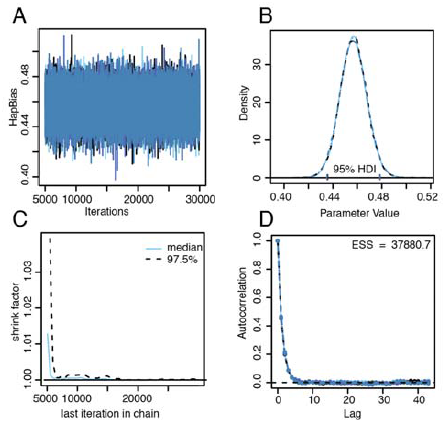
Diagnosis plots of MCMC for type β SNP in family one. (A) The trace plot shows the sampled HapBias value of four independent chains in MCMC simulations. (B) HapBias density distributions of four chains. The limits of 95% HDI are labeled. (C) The trend of shrink factor along increased iteration in simulations, median and 95% quantile values are showed. (D) Autocorrelation calculated with various lags for four chains. Total ESS is showed in the plot. Lag, the number of iterations between the chain and the superimposed copy.

Enough simulation steps are critical for accurately representing the parameter distribution. The autocorrelation plot, calculated as the average correlation between data and k steps ahead data, provides hits on level of redundancy of simulations. Based on the value of autocorrelation, we further calculated the effective sample size (ESS), which summarizes how many completely non-autocorrelated steps in our simulation.^20^ Heuristically, an ESS of 10,000 is recommended for many MCMC simulations.^13^ In our results, we could see autocorrelation value gets close to 0 very quickly for k great than 6, while 37881 ESS were achieved (Fig. 4D, ESS is 34887 in Supplementary Fig. 2D). Thus, the simulated posterior distributions were highly accurate for tested data sets.

### Calculation the limits of the region of practical equivalence

ROPE, which is alternatively named “indifference zone” or "range of equivalence”, means the parameter values within this range are considered to be practically equivalent for the null value^15^. Small difference between null value and estimated value are most likely caused by sampling variation or data error, rather than true difference. The null hypothesis is that HapI and HapII are equally presented, i.e., HapBias is 0.5. For the data generated by next generation sequencing methods, we assumed that the ROPE limits are mainly influenced by sequencing errors,^21^ while higher sequencing error rates lead to wider ROPE. Currently the error rate of next-generation sequencing platform is around 10^-2^,^21,22^ and an error rate of 0.303% has been reported in one NIPD study.^8^ A higher error rate will gave a wider ROPE, leading to the acceptance of null hypothesis. We applied a ROPE of (0.49, 0.51) in the tested data sets, reflecting an error rate of 2% (see Methods). Interestingly, even with this rather conservative ROPE limits, we still successfully inferred the allele inheritance in the real data sets, suggesting the sensitivity of the Bayesian approach.

### Simulation

To further explore the performance of the Bayesian hierarchical model, we generated the high-throughput sequencing data *in silico*. Briefly, we sampled the SNPs counts for HapI and HapII according to the proportion of fetus DNA from a pool of reads, with 500 and 1000 SNPs used, respectively (see Methods). In the condition of 3% fetus DNA, the Bayesian approach correctly detected the maternal allele inherited in all samples using 1000 SNPs, while the detection rate reduced to 76% using 500 SNPs, indicating the benefits of larger data (Supplementary Table 1). SPRT could correctly deduce the maternal allele inheritance in 80% of simulated samples with as low as 0.8% fetus DNA using 1000 SNPs, showing a higher sensitivity (Supplementary Table 2). Taken together, both Bayesian approach and SPRT have great sensitivity in deducing the true maternal allele inheritance.

We next asked how robust these methods are in the existence of measurement errors. The fetus DNA proportion is a key parameter in the SPRT approach as it affects the low and up-boundary of the test statistic under null hypothesis. However, the fetus DNA proportion might not be accurately estimated. We simulated data of 500 SNPs in the conditions that the true fetus DNA proportion was set to be 0% or 10%, and the measured fetus DNA proportion varied from 1% to 20%. In the scenarios that the true fetus DNA proportion is 0%, SPRT provided the wrong inference in all tested data sets as it relied on the fetus DNA proportion for the significance test ((Figure 5A). In the condition that the true fetus DNA proportion is 10% but the estimated value is 20%, SPRT inferred 61% of test samples inherited HapII or undetermined inheritance (Supplementary Table 3). On the contrast, the Bayesian approach does not depend on the fetus DNA proportion. When the Bayesian approach was applied to the simulated data with 0% true fetus DNA proportion, the HDI of HapBias posterior distribution fell completely within the ROPE, indicating that the HapBias was equal to 0.5 and there was no fetus DNA in the sequenced data (Fig. 5B).

**Figure 5.**
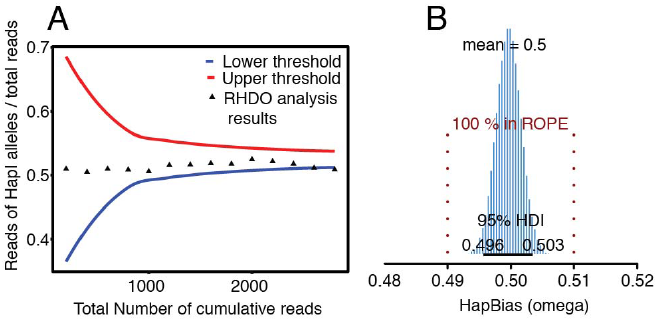
Compare the performance of SPRT and Bayesian model on erroneous data. (A) The SPRT classification incorrectly shows that HapII is inherited in fetus even there is no fetus DNA in sample. (B) Bayesian hierarchical model correctly shows that the true HapBias is equal to 0.5, indicating there is no fetus DNA in samples

### Discussion

Although several methods are available to analyze the NIPD sequencing data, the inference the inheritance of heterogeneous maternal alleles in fetus genome is still a challenging task.^23^ Haplotypes based approaches which uses the imbalance representation of SNPs introduced by fetus DNA provide convincing results^9,12,24^. However, the performance of these methods relies on the value of fetus DNA proportion, which might be not feasible to accurately estimate. We introduced the Bayesian hierarchical model under the framework of RHDO and is independent of fetus DNA proportion in inferring the inherited maternal alleles in fetus genome. To test the null hypothesis that the haplotype frequency is equal to 0.5, we established the posterior distribution of HapBias (proportion of HapI) based on high-throughput sequencing data of individual SNPs, and compared the HDI of HapBias distribution with predefined ROPE to determine which allele is inherited in fetus genome. This method correctly deduced maternal allele inheritances on experimental data, and the results are summarized as intuitive graphs, together with various diagnoses plots.

We evaluated the performance of the Bayesian approach and another widely used approach, SPRT, on simulated data. Both methods could successfully identify the inherited alleles in all samples containing as low as 3% fetus DNA, and SPRT performs well even in samples with 0.8% fetus DNA. Moreover, we found that the Bayesian approach is more robust in conditions where the fetus DNA proportion is not correctly measured. In the extreme case where there is no fetus DNA in the plasma but incorrectly recorded, SPRT provided misleading test results. In contrast, the Bayesian model showed that the HDI of HapBias posterior distribution falls completely within ROPE, corresponding to the hypothesis that there is no fetus DNA in the plasma. Taken together, the Bayesian hierarchical model is a robust approach in deducing the allele inheritance in NIPD.

## Acknowledgement

We thank Dr. Y M Dennis Lo and Dr. JIANG Peiyong for kindly providing the SRPT codes.

## REFERENCES

1. Lo YD, Corbetta N, Chamberlain PF, Rai V, Sargent IL, Redman CW, et al. Presence of fetal DNA in maternal plasma and serum. The Lancet. 1997;350(9076):485–487.

2. Lun FMF, Chiu RWK, Allen Chan KC, Yeung Leung T, Kin Lau T, Dennis Lo YM. Microfluidics Digital PCR Reveals a Higher than Expected Fraction of Fetal DNA in Maternal Plasma. Clin Chem. 2008 Aug 14;54(10):1664–72.

3. Galbiati S, Smid M, Gambini D, Ferrari A, Restagno G, Viora E, et al. Fetal DNA detection in maternal plasma throughout gestation. Hum Genet. 2005 Jul;117(2-3):243–8.

4. Devaney SA, Palomaki GE, Scott JA, Bianchi DW. Noninvasive Fetal Sex Determination Using Cell-Free Fetal DNA: A Systematic Review and Meta-analysis. JAMA [Internet]. 2011 Aug 10 [cited 2016 Apr 18];306(6). Available from: http://jama.jamanetwork.com/article.aspx?doi=10.1001/jama.2011.1114

5. Zhao C, Tynan J, Ehrich M, Hannum G, McCullough R, Saldivar J-S, et al. Detection of fetal subchromosomal abnormalities by sequencing circulating cell-free DNA from maternal plasma. Clin Chem. 2015 Apr;61(4):608–616.

6. Yin A, Peng C, Zhao X, Caughey BA, Yang J, Liu J, et al. Noninvasive detection of fetal subchromosomal abnormalities by semiconductor sequencing of maternal plasma DNA. PNAS. 2015 Nov;112(47): 14670–14675.

7. Ferres MA, Hui L, Bianchi DW. Antenatal Noninvasive DNA Testing: Clinical Experience and Impact. Amer J Perinatol. 2014 Jun 24;31(7):577–82.

8. Lo YMD, Chan KCA, Sun H, Chen EZ, Jiang P, Lun FMF, et al. Maternal plasma DNA sequencing reveals the genome-wide genetic and mutational profile of the fetus. Sci Transl Med. 2010 Dec;2(61):61ra91.

9. Lam KWG, Jiang P, Liao GJW, Chan KCA, Leung TY, Chiu RWK, et al. Noninvasive Prenatal Diagnosis of Monogenic Diseases by Targeted Massively Parallel Sequencing of Maternal Plasma: Application to β-Thalassemia. Clin Chem. 2012 Sep;58(10):1467–1475.

10. New MI, Tong YK, Yuen T, Jiang P, Pina C, Chan KCA, et al. Noninvasive Prenatal Diagnosis of Congenital Adrenal Hyperplasia Using Cell-Free Fetal DNA in Maternal Plasma. J Clin Endocrinol Metab. 2014 Jun;99(6):E1022–E1030.

11. Fan HC, Gu W, Wang J, Blumenfeld YJ, El-Sayed YY, Quake SR. Non-invasive prenatal measurement of the fetal genome. Nature. 2012 Jul 4;487(7407):320–4.

12. Kitzman JO, Snyder MW, Ventura M, Lewis AP, Qiu R, Simmons LE, et al. Noninvasive whole-genome sequencing of a human fetus. Sci Transl Med. 2012 Jun;4(137):137ra76.

13. Kruschke JK. Doing Bayesian data analysis: a tutorial with R, JAGS, and Stan. Edition 2. Boston: Academic Press; 2015. 759 p.

14. Hobbs BP, Carlin BP. Practical Bayesian Design and Analysis for Drug and Device Clinical Trials. J Biopharm Stat. 2007 Dec 10;18(1):54–80.

15. Freedman LS, Spiegelhalter DJ. The Assessment of the Subjective Opinion and its Use in Relation to Stopping Rules for Clinical Trials. The Statistician. 1983Mar;32(1/2):153.

16. Berry SM. Bayesian adaptive methods for clinical trials [Internet]. Boca Raton: CRC Press; 2011.

17. Schafer JL. Analysis of incomplete multivariate data. CRC Press; 2000.

18. Enders CK. Applied missing data analysis. New York: Guilford Press; 2010.

19. Brooks SP, Gelman A. General Methods for Monitoring Convergence of Iterative Simulations. J Comput Graph Stat. 1998 Dec;7(4):434–55.

20. Kass RE, Carlin BP, Gelman A, Neal RM. Markov Chain Monte Carlo in Practice: A Roundtable Discussion. Am Stat. 1998 May;52(2):93–100.

21. Fox EJ, Reid-Bayliss KS, Emond MJ, Loeb LA. Accuracy of next generation sequencing platforms. Gener Seq Appl. 2014.

22. Victoria X, Blades N, Ding J, Sultana R. Estimation of Sequencing Error Rates in Short Reads. BMC Bioinformatics, 2012.

23. Chan LL, Jiang P. Bioinformatics analysis of circulating cell-free DNA sequencing data. Clin Biochem. 2015 Oct;48(15):962–75.

24. Lun FM, Tsui NB, Chan KA, Leung TY, Lau TK, Charoenkwan P, et al. Noninvasive prenatal diagnosis of monogenic diseases by digital size selection and relative mutation dosage on DNA in maternal plasma. Proc Natl Acad Sci. 2008;105(50):19920–19925.

